# SARS-CoV-2 ORF9c Is a Membrane-Associated Protein that Suppresses Antiviral Responses in Cells

**DOI:** 10.1101/2020.08.18.256776

**Authors:** Ana Dominguez Andres, Yongmei Feng, Alexandre Rosa Campos, Jun Yin, Chih-Cheng Yang, Brian James, Rabi Murad, Hyungsoo Kim, Aniruddha J. Deshpande, David E. Gordon, Nevan Krogan, Raffaella Pippa, Ze’ev A. Ronai

## Abstract

Disrupted antiviral immune responses are associated with severe COVID-19, the disease caused by SAR-CoV-2. Here, we show that the 73-amino-acid protein encoded by *ORF9c* of the viral genome contains a putative transmembrane domain, interacts with membrane proteins in multiple cellular compartments, and impairs antiviral processes in a lung epithelial cell line. Proteomic, interactome, and transcriptomic analyses, combined with bioinformatic analysis, revealed that expression of only this highly unstable small viral protein impaired interferon signaling, antigen presentation, and complement signaling, while inducing IL-6 signaling. Furthermore, we showed that interfering with ORF9c degradation by either proteasome inhibition or inhibition of the ATPase VCP blunted the effects of ORF9c. Our study indicated that ORF9c enables immune evasion and coordinates cellular changes essential for the SARS-CoV-2 life cycle.

**One-sentence summary:** SARS-CoV-2 ORF9c is the first human coronavirus protein localized to membrane, suppressing antiviral response, resembling full viral infection.

## Introduction

SARS-CoV-2 is an enveloped, positive-sense single strand 29.9 kb RNA virus (*1, 2*) that causes severe respiratory disease in humans (COVID-19). This coronavirus was first identified in Wuhan, China, at the end of 2019 (*3*). Due to its easy human-to-human transmission and the lack of effective antiviral therapy, COVID-19 has caused a pandemic with more than 19 million cases and over 740,000 deaths worldwide (https://covid19.who.int). Mechanistically, the host protein ACE2 serves as the viral receptor and host cellular proteases, such as TMPRSS2, play key roles in SARS-CoV-2 entry into host cells (*4–7*). *ACE2* expression is high in alveolar epithelial cells (*8*), making the lung a highly vulnerable target for the virus.

SARS-CoV-2 infection causes a wide range of disease, from asymptomatic to mild disease to severe disease that can lead to death (*9*). SARS-CoV-2 is most similar to the coronaviruses SARS-CoV and MERS-CoV (*10, 11*). However, neither of those became a global pandemic. Current therapies are primarily palliative and supportive (*9*). More than 2000 clinical trials are currently in progress worldwide (*12*) (https://clinicaltrials.gov/ct2/who_table). Without effective vaccines or treatments, there is an urgent need to understand the pathology of SARS-CoV-2 infection, the roles of each of the 29 proteins encoded within the viral genome in the life cycle, virulence, and pathogenicity of the virus, and identify strategies for intervention or treatment.

Various therapeutic and vaccine strategies target viral entry mechanisms, such as vaccines or antibodies targeting on the Spike (S) protein (*13–16*); others target viral replication or assembly processes, such as the antiviral drug remdesivir, which interferes with RNA replication and has emerged as superior to placebo in shortening recovery time in adults (*17*). Another strategy for treatment is interfere with viral immune evasion mechanisms and thus enable the body’s natural antiviral responses to be more effective at clearing the virus. Indeed, investigation of mechanisms of immune evasion by SARS-CoV-2 is an active area of translational research with immune evasion properties discovered for nonstructural protein 1 (Nsp1) (*18*).

The SARS-CoV-2 genome contains 15 open reading frames (ORFs), which encode 29 viral proteins (*19–21*). ORF1a and ORF1ab encode polyproteins that are cleaved into 16 nonstructural proteins (Nsp1 – Nsp16) that comprise the replicase-transcriptase complex. Spike (S) is encoded by ORF2, envelope (E) by ORF4, membrane (M) by ORF5, and nucleocapsid (N) by ORF9. An additional 9 ORFs encode “accessory” proteins: ORF3a, ORF3b, ORF6, ORF7a, ORF7b, ORF8, ORF9b, ORF9c, and ORF10.

Various studies have investigated the functions of the virally encoded proteins by performing interactome analysis in cells expressing individual viral proteins (*19*) or by evaluating the proteomic or transcriptomic changes associated with either viral infection (*22–27*). Others have used computational approaches to investigate protein-protein interactions between SARS-CoV-2 viral proteins and host proteins (*28*). The interactome and proteome studies identified cellular processes affected by SARS-CoV-2 infection or specific viral proteins, notably innate immune signaling (*19, 20, 23, 28–30*), ubiquitin ligase activities (*19, 20, 23, 28–30*), p38 mitogen-activated protein kinase (MAPK) signaling (*19, 20, 23, 28–30*). The transcriptomic studies identified interferon signaling (*24*), cell death (*27*), interleukin 1 (IL-1), IL-6, and chemokine signaling (*22*).

Given the intense interest in catalytically active CoV-2 proteins (*31–34*), we examined the less-studied group of ORFs encoding accessory proteins, which are largely thought to maintain viral structural organization in replication organelles and within the viral particle (*35, 36*). Here, we showed that expression of only ORF9c is sufficient to alter cellular networks in a manner that resembles full SARS-CoV-2 virus infection.

## Results

### SARS-CoV-2 ORF9c encodes an unstable protein with a putative transmembrane domain

Seven of the 29 CoV-2 proteins are ORFs that lack catalytic activity and, in some cases, lack a known function (*19*). Each of these were tagged with Strep at the N terminus and expressed them individually in the lung cancer epithelial cell line A549 in the presence or absence of the proteasome inhibitor MG132 (Fig. 1A). The protein encoded by ORF9c was particularly unstable, with a profound increase in abundance evident in MG132-treated compared to that in vehicle-treated A549 cells (Fig 1A).

**Figure 1.**
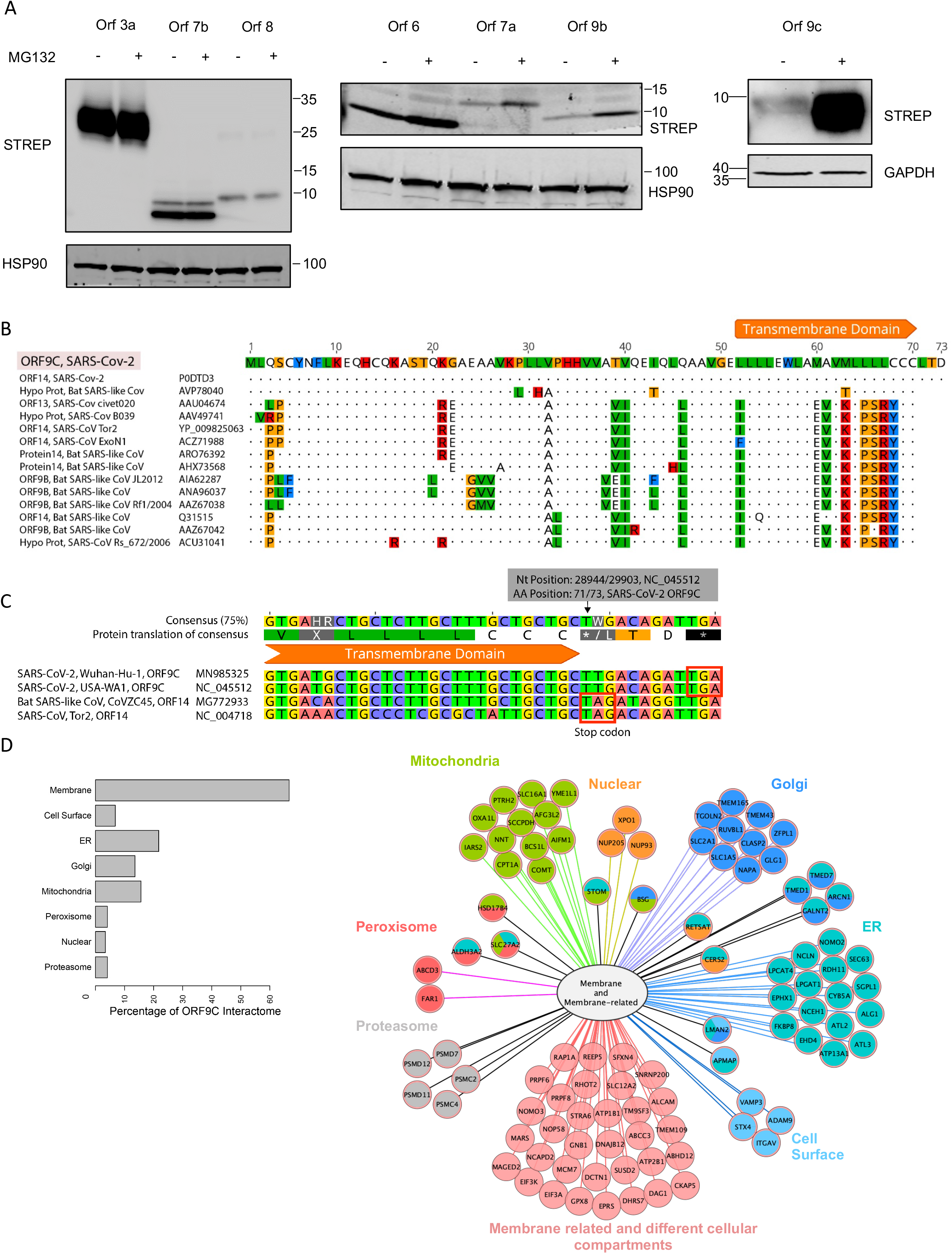
The SARS-CoV-2 ORF9c interactome in A549 cells. **A.** Stability of accessory proteins encoded by SARS-CoV-2 ORFs in A549 cells. Proteins were prepared 24 h after transfection of lung cancer A549 cells with the indicated Strep-tagged ORF constructs in the presence or absence of MG132 (10 μM added 4 h before harvest). Proteins were detected Western blotting with antibodies to Strep. **B**. Sequence alignment of SARS-CoV-2 ORF9c to related orthologs. Dots indicate identity. Amino acid color is displayed using Clustal scheme as hydrophobic (I, L, M, V) as green, aromatic (F, W, Y) as blue, positive charge (K, R, H) as red, proline/glycine and some hydrophilic polar amino acids (G,P,S,T) as orange. **C.** Alignment of the gene sequence showing the position of the stop codon in ORF14 orthologs and 3 codon extension in ORF9c. **D**. Interactome of ORF9c is based on LC-MS/MS of ORF9c-interacting proteins immunoprecipitated from A549 24 h after transfection. Left: Number of ORF9C-interacting proteins according to their cell compartment (from Gene Ontology). Total values exceed 100% because some proteins are assigned / located in more than one compartment. Right: Protein subcellular localization map for the portion of the ORF9c interactome that is either a membrane protein or a membrane-related protein (as defined by Gene Ontology).

ORF9c is present in previously characterized strains of SARS-CoV (*37*), a conservation suggesting a function in coronavirus pathogenesis. Phylogenetic analysis and alignment of the protein sequences showed that mutations are present in ORF9c among different coronavirus strains with bat SARS-like coronavirus ORF14 as the closest ortholog sharing 94% sequence identity and only 77% identity with ORF14 of SARS-CoV (Fig. 1B, fig. S1A). TMHMM analysis (*38*) of SARS-CoV-2 ORF9c predicted a transmembrane sequence in the C-terminal domain, a motif not present in SARS-CoV-1 (or other human coronaviruses) ORF9c sequence (Fig. 1B, fig. S1B). Additionally, a single nucleotide mutation in SARS-CoV-2 ORF9c altered a termination codon, enabling the reading frame to extend by 3 amino acids (Fig. 1C).

### SARS-CoV-2 ORF9c interactome includes membrane-associated proteins distributed throughout multiple cellular compartments

We assessed potential functions of ORF9c by performing transcriptome, interactome, and proteome analysis of A549 lung cancer cells transfected with ORF9c tagged at the N terminus with 2 copies of the Strep tag (*19*). To map the ORF9c interactome, we conducted liquid chromatography tandem mass spectrometry (LC-MS/MS) of 2xStrep-tagged ORF9c compared with control 2xStrep-tagged GFP) immunoprecipitated from A549 cells 24 h after transfection. ORF9c interactome analysis revealed that most interacting proteins were classified as membrane proteins (Fig. 1D, table *S1*) according to Gene Ontology Cellular Component. As a protein with a transmembrane domain, this was not surprising. However, we were surprised to find that the ORF9c interactome was distributed throughout the membrane-bound organelles (Fig. 1D), including >30 proteins in the protein biosynthesis and transport systems of the endoplasmic reticulum (ER) and Golgi, >15 proteins in the mitochondria, and >30 other membrane-related proteins. Given the instability of ORF9c, we were not surprised to identify a group of membrane-associated proteins that function with the proteasome. Comparison between ORF9c and ORF10, both expressed using the Strep-tagged vector, confirmed that enrichment of membranal proteins as part of the interactome was selectively seen for the ORF9c (fig. S1C).

### Proteome analysis shows ORF9c downregulates proteins involved in interferon signaling and antigen processing

We conducted both label free quantification (LFQ) and tandem mass tag (TMT) mass spectrometry analysis to identify changes in the cellular proteome in A549 cells expressing ORF9c. We compared the proteomic changes associated with ORF9c expression in the presence or absence of proteasomal inhibition with MG132, using DMSO as the vehicle control for comparison in each set. Principle component analysis (PCA) revealed that ORF9c contributed to major variance in all data sets (fig. S2A, S2B). Pairwise comparisons between ORF9c and control untransfected samples identified differentially expressed proteins in both the DMSO and MG132 groups (Fig 2A). In both the DMSO and MG132 datasets, most changes induced by ORF9c were a reduction in protein abundance (downregulation) (Fig. 2A, table S1).

**Figure 2.**
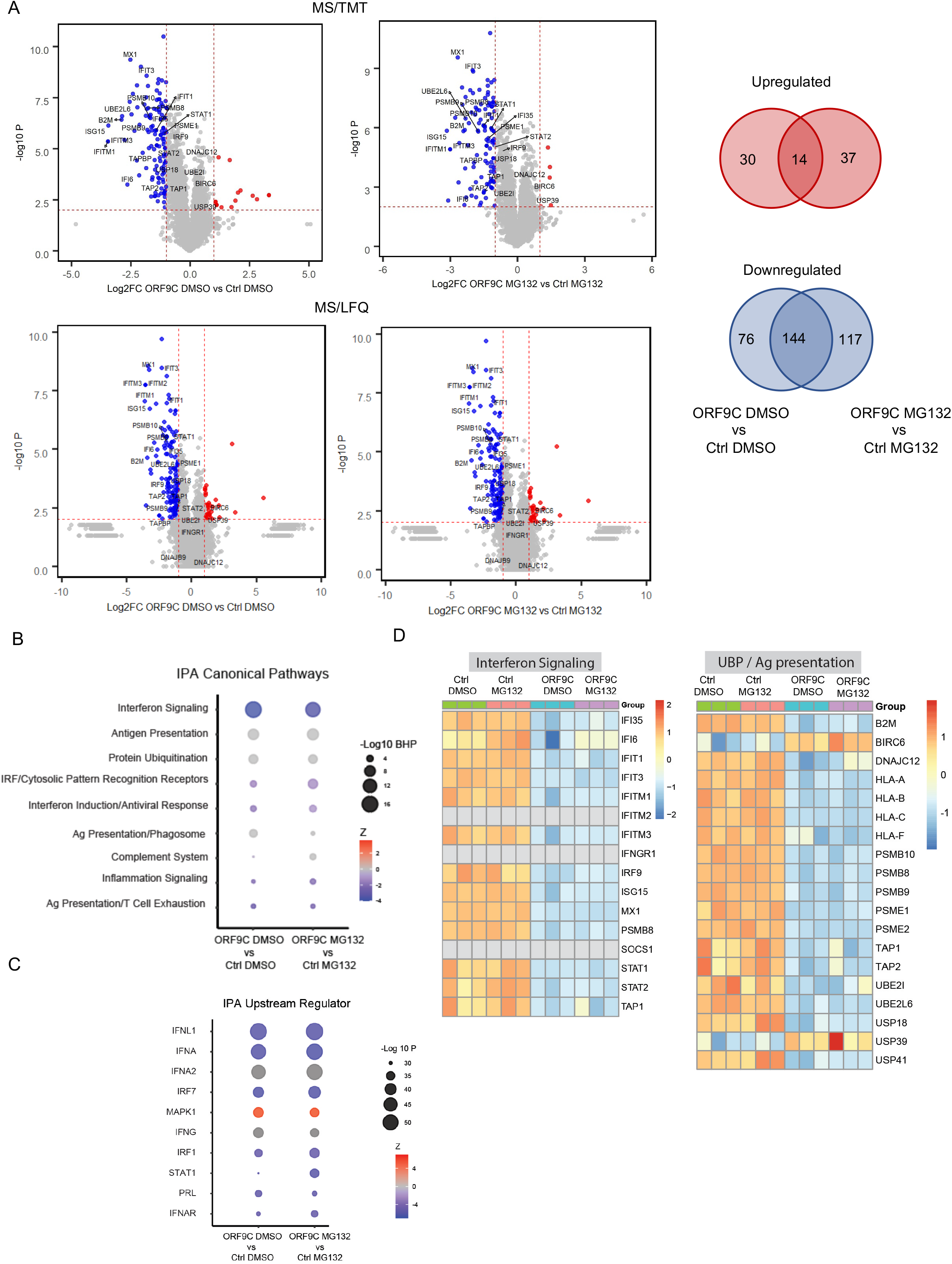
Proteomic profiling of A549 cells expressing SARS-CoV-2 ORF9c. **A.** Proteomic profiling of A549 cells infected with Strep-tagged SARS-CoV-2 ORF9c compared Strep-tagged GFP as a control. Left: Volcano plots show differentially regulated proteins from MS/TMT or MS/LFQ analysis. Red dots, upregulated proteins; blue dots, downregulated proteins. Right: Venn diagrams show intersection of up- or down-regulated proteins between ORF9c-infected A549 versus control cells cultured in DMSO vehicle or MG132 from both MS/TMT and MS/LFQ analysis. **B**. Differentially regulated proteins identified in both MS/TMT and MS/LFQ were subjected to IPA “canonical pathway” analyses. Shown are pathways enriched based on differentially regulated proteins (comparing ORF9c with control). Dot size is scaled to −log_10_ Bonferroni Hochberg-corrected P (BHP) value. Dot color represents a gradient of *Z* scores with score predicting pathway activity: A positive score indicates activation; a negative score indicates inhibition. **C**. IPA “upstream regulator” analyses for differentially regulated proteins (comparing ORF9c with control). Dot size is scaled to −log_10_ P value. Dot color represents a gradient of Z scores with score predicting change in abundance: A positive score indicates increase; a negative score indicates decrease. **D**. Heatmap representation of differentially regulated proteins classified as interferon signaling or as protein ubiquitination-proteasome (UBP) and antigen (Ag) presentation. Data are from MS/TMT analysis. Color bars present normalized abundance values.

Downregulated proteins identified using both approaches consistently showed ~60% overlap, while no overlap were identified among the upregulated proteins. Thus, to maximize the discovery of ORF9c dysregulated proteins, results from both technologies were combined. Including the differentially regulated proteins identified by both the TMT and LFQ analysis revealed 14 proteins were upregulated in common by ORF9c expression in the presence or absence of the proteasome inhibitor and 144 proteins were downregulated in common (Fig. 2B). Using the downregulated proteins and upregulated proteins identified for either the DMSO or MG132 condition separately, we performed Ingenuity Pathway Analysis (IPA) to assess signaling pathways deregulated in ORF9c-expressing cells. In both the DMSO and MG132 condition, interferon (IFN) signaling exhibited the greatest difference, both in terms of the intensity of the downregulation and the number of proteins significantly associated with this pathway, in response to ORF9c (Fig. 2C, table S1). Other pathways affected by ORF9c and of particular importance to virulence were antigen presentation and innate immune response pathways, such as IRF/cytosolic pattern recognition receptors. We further examined potential upstream regulators of these pathways using Ingenuity Pathway Analysis for proteins that exhibited a change in abundance in the ORF9c-expressing cells. This analysis revealed that several components of the IFN machinery [interferons (IFNL1, IFNA), interferon responsive transcription factors (IRF7, IRF1), and an interferon receptor (INFAR)] were reduced, consistent with the impaired IFN signaling, and an increase in MAPK1 (also known as ERK2) abundance (Fig. 2C, table S1).

To assess if there were notable differences in the intensity of the changes in protein abundance in response to proteasome inhibition, we calculated relative changes in protein abundance between control and ORF9c-expressing cells from both the DMSO and MG132 conditions for proteins associated with IFN signaling or the ubiquitin proteasome (UBP) system and antigen presentation (Fig. 2D). The intensity of the changes was similar in the presence or absence of MG132, suggesting even small amounts of the unstable ORF9c are sufficient to induce cellular changes including those that contribute to immune evasion.

Consistent with the IPA-based analysis (Fig. 2B), IFN signaling components, including IFI35, multiple IFIT proteins, IRF9, ISG15, MX1, PSMB8, and STAT proteins, were downregulated in ORF9c-expressing cells in both the presence and absence of MG132 (Fig. 2D, left). Indicative of a decrease in antigen presentation capacity, multiple proteins involved in this process were decreased, including proteins involved in antigen loading and display [HLA proteins, β2M, and antigen transporters (TAP1 and TAP2)] and proteins involved in UBP [ubiquitin-conjugating enzymes UBE2I and UBE2L6), deubiquitination enzymes (USP18 and UPS41), and proteasome components (PSMB and PSME proteins)] (Fig 2D, right).

These changes in the proteome indicated that the expression of only ORF9c, even in the absence of proteasomal inhibition to stabilize this protein, is sufficient to elicit effective inhibition of IFN, immune recognition, and UBP components at the protein level. Such a response suggested that ORF9c contributes to immune evasion of SARS-CoV-2.

### ORF9c attenuates transcription of immune and stress signaling pathways

We assessed transcriptional changes elicited by ORF9c expression in A549 cells using RNA-seq analysis. PCA showed that changes induced by ORF9c in both DMSO- and MG132-treated cells cluster in distinct experimental groups (Fig. S2C). In contrast to the proteomic results that revealed predominant downregulation of proteins following ORF9c expression, RNA-seq analysis showed a similar number of transcripts were increased or decreased in the presence or absence of MG132 (Fig. 3A, table S2). Additionally, the number of differentially regulated transcripts was higher than that for the differentially regulated proteins.

**Figure 3.**
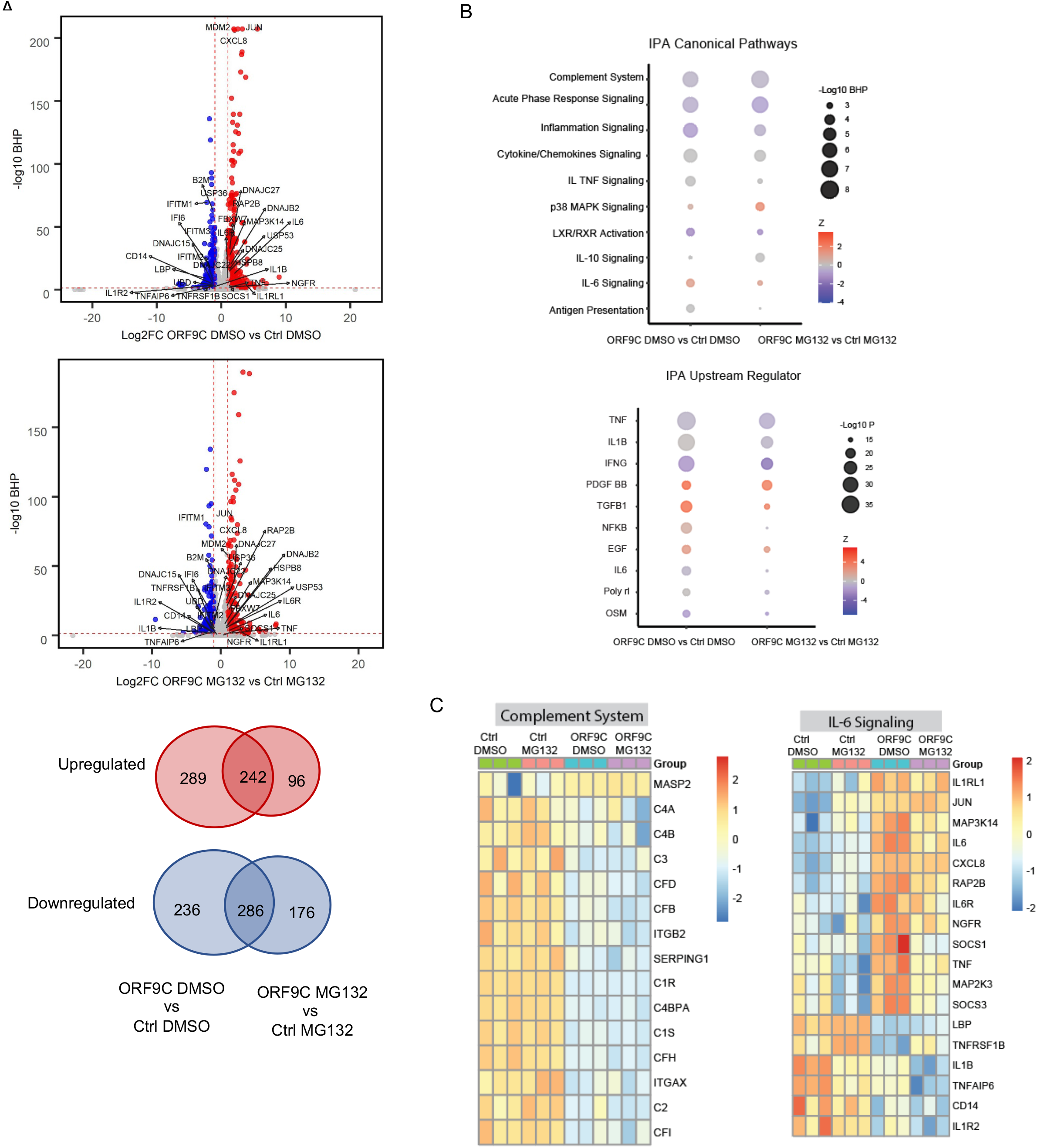
Transcriptomic profiling of A549 cells expressing SARS-CoV-2 ORF9c. **A**. Volcano plots show differentially abundant transcripts based on RNA-Seq analyses. Red dots, up-regulation; blue dots, downregulation. Venn diagrams depict intersection of up- or down-regulated genes between ORF9c-infected A549 cells versus control cells. **B**. IPA “canonical pathway” and “upstream regulator” analyses of differentially expressed genes from ORF9c compared to control cells. Dot size is scaled to −log_10_ BH-corrected P value (BHP) or −log_10_ P value. Dot color represents a Z score gradient with score predicting activity or abundance: A positive score indicates activation or increase, and a negative score indicates inhibition or decrease. **C.** Heatmap representation of differentially expressed genes associated with the complement system or IL-6 signaling, based on RNA-Seq. Color bars present normalized expression values.

Using IPA, we identified the pathways significantly enriched in differentially regulated transcripts. The same set of pathways were identified in the DMSO and MG132 conditions, and similar to the proteomic results, most related to immune signaling (Fig. 3B, upper, table S2). However, many of the specific pathways were different from those identified at proteomic level. At the transcriptional level, we detected the greatest effects on the complement system and several pathways involved in inflammatory signaling. Thus, some components of antigen presentation and immune signaling pathways showed comparable changes at the protein and mRNA levels; other changes elicited by ORF9c expression were unique to the transcriptional level, such as induction of IL-6 signaling and p38 MAPK signaling, or the protein level, such as impairment of IFN signaling.

We analyzed the ORF9c-regulated transcripts for those encoding upstream regulators of the pathways altered at the transcriptional level by ORF9c. This analysis identified the classic immune modulators tumor necrosis factor (TNF), IL-1B, IFNγ, transforming growth factor β (TGFβ), and NF-κB signaling components (Fig. 3B, lower, table S2).

We evaluated transcripts associated with the complement system or IL-6 signaling in detail. In both the presence and absence of proteasome inhibition, complement system transcripts were mostly downregulated by ORF9c expression in A549 cells (Fig. 3C, left). For IL-6 signaling, we found some differences between the MG132 and DMSO conditions (Fig. 3C, right). For several transcripts the intensity of the upregulation was greater in the absence of proteasome inhibition (MAP3K14 and MAP2K3, IL6 and IL6R, SOCS1 and SOCS3); for others the presence of the proteasome inhibitor resulted in a greater reduction in transcript abundance (IL1B, TNFAIP6, CD14, IL1R2). Thus, these results suggested that ORF9c had a dose-dependent effect on some transcripts.

### Common proteome and transcriptome changes induced by ORF9c

We combined the results of the proteomic and transcriptomic (Fig. 4A), which revealed a small set of commonly upregulated or downregulated genes by ORF9c at both the transcription and protein levels (Fig. 4B). We performed IPA canonical pathway analysis and found commonly altered pathways at both the transcript and protein levels (Fig. 4C, table S1, S2). The direction of regulation (increased or decreased activity) was consistent between the transcripts and proteins. However, the number of components significantly enriched in most of the pathways differed between the protein and transcript levels.

**Figure 4.**
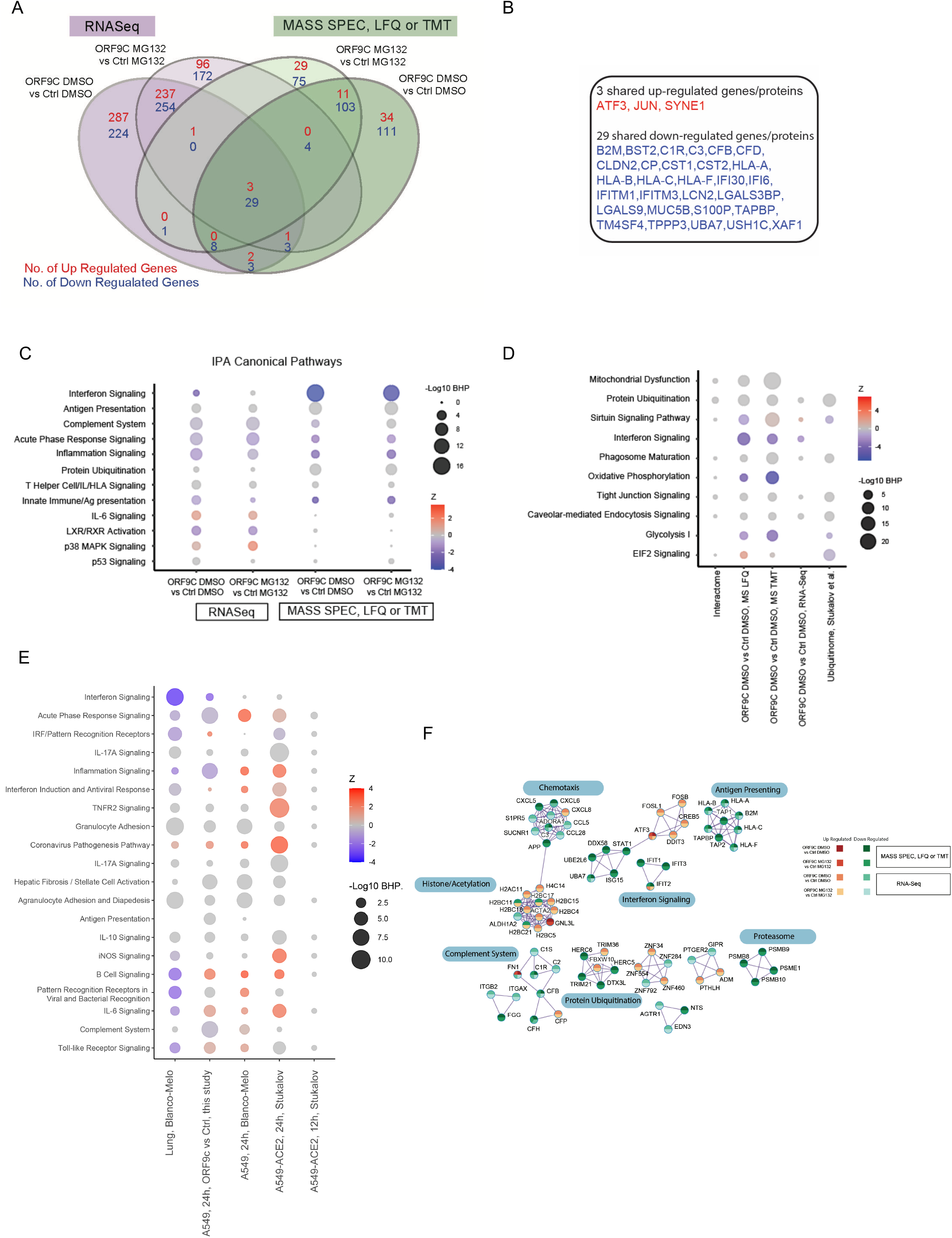
Comparison of transcriptomic and proteomic profiles of A549 cells expressing SARS-CoV-2 ORF9c. **A**. Venn diagram depicting intersection of up- or down-regulated genes or proteins between ORF9c-expressing A549 versus control cells, both under control (DMSO) or MG132 conditions. **B**. List of the 3 common upregulated genes or protein and 29 downregulated genes or proteins common to all data sets. **C.** IPA “canonical pathway” analyses of differentially expressed genes or proteins from ORF9c-expressing compared to control cells. Dot color represents a gradient of Z scores with score predicting pathway activity: A positive score indicates activation; a negative score indicates inhibition. **D**. IPA-based comparison of pathways enriched based on different analyses indicated. Comparison is based on ORF9c interactome, proteome and transcriptome data from this study and that of a published data set (*39*). Dot size and color as described in C. **E**. IPA-based comparison of pathways enriched based on different transcriptome analyses from this study and the indicated published data sets from SARS-CoV-2–infected cells (*22, 39*). Lung, Blanco-Melo, data are SARS-CoV-2 infected primary lung epithelial cells; A549, Blanco-Melo, data are from A459 cells and infected with SARS-CoV-2; A549-ACE2, 24h, Sukalov and A549-ACE2, 12h, Sukalov, data are from A459 cells expressing ACE2 and infected with SARS-CoV-2 12 hours and 24 hours after infection. Dot size and color as described in C. **F**. Protein-protein interaction network of the ORF9c-dysregulated transcriptome and proteome. The protein-protein interaction network was defined by the Molecular Complex Detection (MCODE) algorithm based on known protein physical interactions. Protein color represents direction of changes in ORF9c versus control cells grown in DMSO or MG132.

We compared our findings at the transcriptome, proteome, and interactome levels with those reported by Stukalov et al. (*39*) for proteins with altered ubiquitination (ubiquitinome) in response to SARS-CoV-2 infection of A549 cells. Within the top 10 enriched IPA canonical pathways, we noticed enrichment across all 5 protein ubiquitination data sets, sirtuin signaling, phagosome maturation, tight junction signaling, and caveolar-mediate endocytosis (Fig. 4D, table S3). Seven of the 10 were common between the ubiquitinome data (*39*) and our proteomic analyses by LFQ or TMT mass spectrometry.

We also compared the pathway enrichment across our transcriptomic data and those from Blanco-Melo *et al*. (*22*) reporting transcriptomic changes upon SARS-CoV-2 infection in human primary epithelial cells or ACE-expressing A549 cells and with those from Stukalov *et al*. (*39*) reporting transcriptomic changes in ACE2-expressing A549 cells 12 and 24 hours after infection (Fig. 4E, table S3). The changes in cellular pathways that we observed at the transcriptional level in A549 cells expressing only ORF9c were remarkably similar to those observed in SARS-CoV-2-infected primary lung cells with a few exceptions. We observed an increase in transcripts associated with B cell signaling and IL-6 signaling in the ORF9c-expressing A549 cells. In contrast, the other studies of ACE2-expressing A549 cells infected with SARS-CoV-2 showed similar pathway responses, but these were different from the infected primary lung epithelial cells. To visualize how the transcript and protein level changes induced by ORF9c were related, we used the Molecular Complex Detection (MCODE) algorithm, which uses physical interactions among components, to construct interaction networks overlaid with the direction of change in either the MG132 or DMSO condition used for mapping changes in transcript and protein levels (Fig. 4A, 4F, table S1, S2). This analysis revealed a complement module, a chemotaxis module, an interferon module, and an antigen presentation module, all of which were mostly downregulated and often coordinately downregulated at both the transcript and protein levels. Two modules appeared primarily induced at the transcript level, a histone acetylation module and a module containing the stress responsive AP1 family members (FOSB, FOSL1, CREB, and ATF3), suggesting that transcriptional repression may underlie some gene expression changes elicited by ORF9c expression.

### ERAD / proteasome inhibitors can partially reverse cellular changes elicited by SARS-CoV-2 ORF9c

In the presence or absence of the proteasome inhibitor, we observed substantial overlap (55 – 65%) in the ORF9c-downregulated proteins (Fig. 2A) and 54 – 72% overlap in ORF9c-upregulated transcripts and 45 – 64% in the downregulated transcripts (Fig. 3A). Thus, despite ORF9c increasing in MG132-treated A459 cells, the persistent changes in MG132-treated cells suggested that even a low amount of ORF9c is sufficient to elicit its cellular effects. However, some transcripts (399) and proteins (97) were downregulated by ORF9c in cells not treated with MG132 and were upregulated in cells treated with MG132 (Fig. 5A).

**Figure 5.**
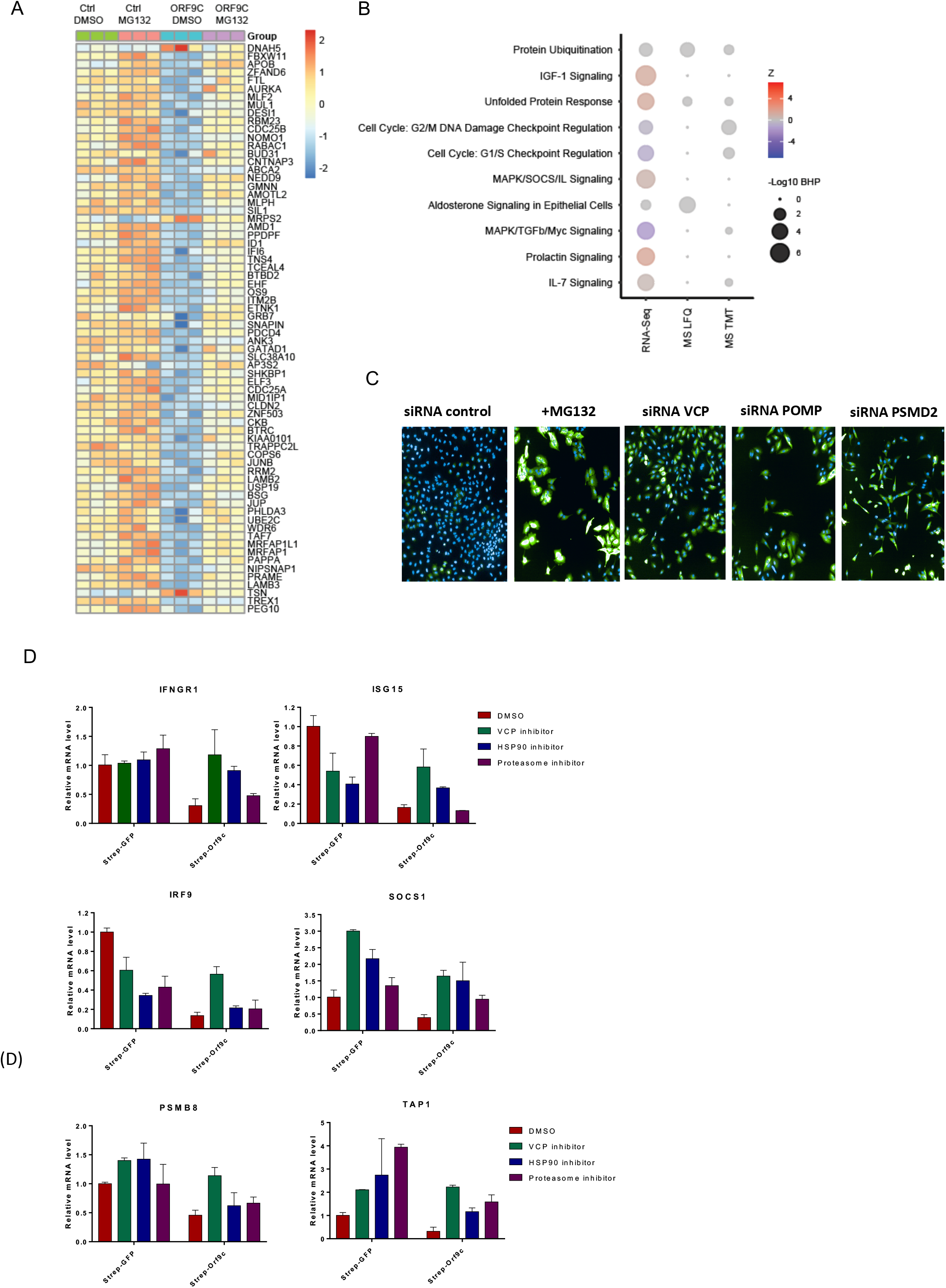
The UBP and UPR control ORF9c stability and partially controls its cellular activity. **A**. Heatmap analysis of the cellular proteins that show opposing direction of regulation in cells expressing ORF9c and exposed to MG132 treatment compared to ORF9c-expressing cells exposed to DMSO. Data are from MS TMT. **B.** IPA “canonical pathway: analysis of cellular pathways enriched following MG132 treatment of indicated treatment groups. **C**. Representative data showing the increase in ORF9c abundance in cells in which the indicated proteins were silenced. Cells exposed to MG132 served as a positive control. **D**. Quantitative RT-PCR analysis of the indicated transcripts in A549 cells stably expressing SARS-CoV-2 ORF9c and exposed to the indicated inhibitors [VCP inhibitor MNS-873 (2 μM), HSP90 inhibitor Geldanamycin (0.1 μM), and proteasome inhibitor Bortezomib (15 nM)] for 24 h. Cells expressing Strep-GFP served as a control for the effects of the inhibitors in the absence of ORF9c.

Pathway analysis on the transcripts and proteins showed discordant regulation in DMSO- or MG132-treated cells in response ORF9c. The most pronounced among all three datasets were components of the UBP and the unfolded protein response (UPR) (Fig. 5B, table S4). Changes in events or pathways associated with the cell cycle was not surprising given the critical role UBP plays in their regulation. We thus hypothesized that both the UBP and UPR were involved in degradation of ORF9c. Finding that UPR signaling was also reversed upon MG132 treatment (Fig 5B) is consistent with the role of UPR signaling in degradation of ORF9c.

To directly assess the importance of UBP and UPR components in ORF9c instability, we performed an siRNA-based screen targeting over 1100 genes that encode components of both machineries in A549 cells stably expressing Strep-tagged ORF9c (Fig. S3). The top hits independently validated as blocking ORF9c degradation were siRNAs targeting VCP [also known as p97, an ATPase involved in export of unfolded proteins from the ER for ER-associated degradation (ERAD) and in ER to Golgi transport] (*40, 41*), the proteasomal subunit PSMD2, and the proteasome maturation factor POMP (*42*), which has been also implicated in ERAD (*43*) and in IFN-induced reorganization of proteasomes into immunoproteasomes (*44*) (Fig. 5C, Table 1).

**Table 1.**
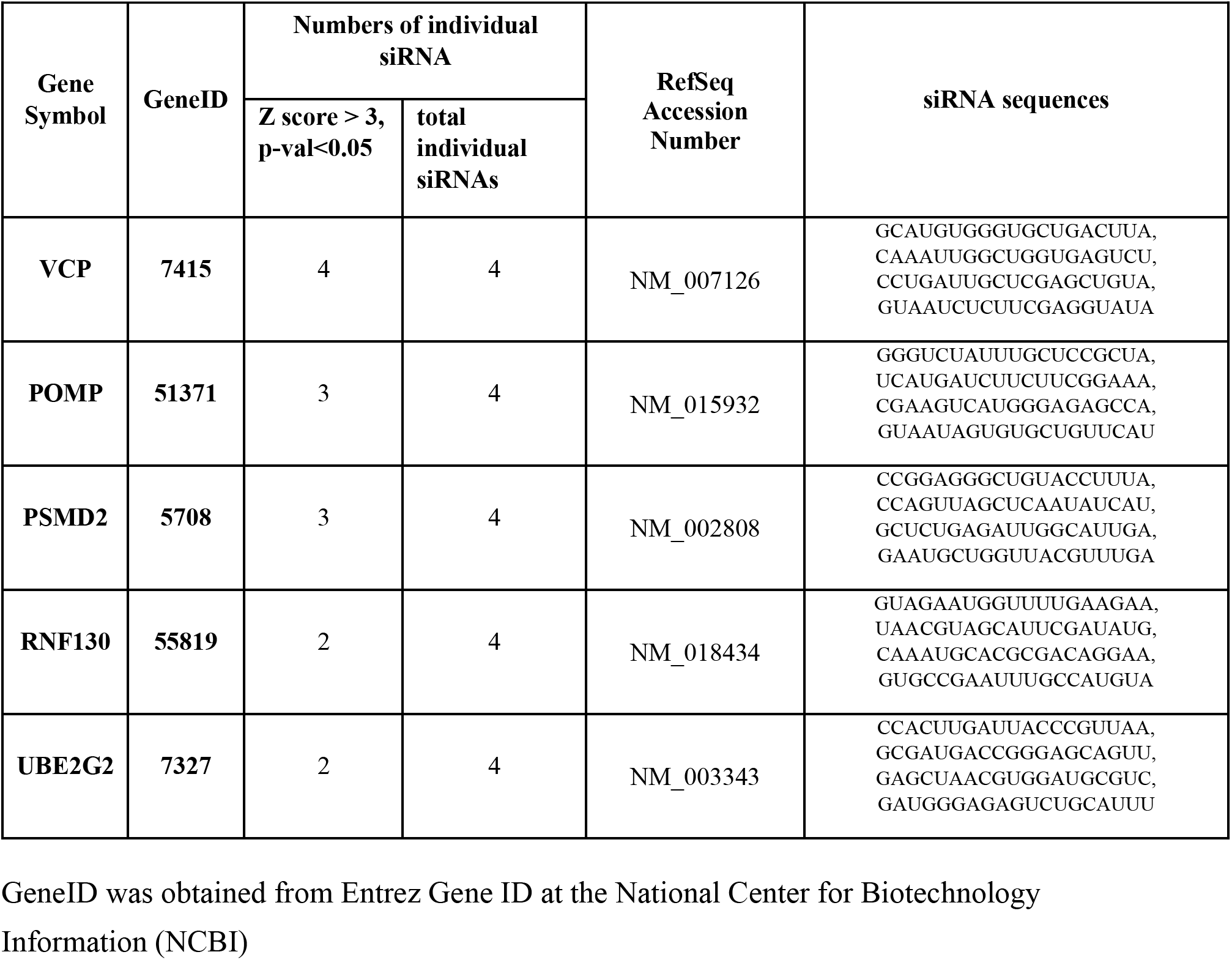
Confirmed siRNA targets that stabilize ORF9c when knocked down in A549 cells.

To assess if interfering with ERAD affected the cellular effects induced by ORF9c, we compared the transcript abundance for 6 genes (*IFNGR1*, *IGS15*, *IRF9*, *SOCS1*, *PSMB8*, *TAP1*) that were downregulated by expression of ORF9c in DMSO-treated cells with their abundance in cells treated with the VCP inhibitor MNS-873, the heat shock protein 90 (HSP90) inhibitor geldanamycin, or the proteasome inhibitor bortezomib. Although the HSP90 inhibitor and the proteasome inhibitor increased transcript abundance for some of the gens tested, VCP inhibition was the most consistently effective at enabling expression of each of these transcripts in the ORF9c-expressing cells (Fig. 5D). These observations suggested that ORF9c ability to attenuate key cellular signaling involved in antiviral responses, including antigen presentation, immune, and IFN pathways, requires the activity of VCP.

## Discussion

The key to our ability to control spread of the SARS-CoV-2 virus is to understand its mechanism of action and how the concerted action of its 29 encoded proteins subvert cellular regulatory networks. One can divide viral “success” into two key phases: infection, which is the ability to enter a given cell type, and multiplication that enables continuous infection through viral replication and packaging, which exploits host cell machineries (*45*). A third aspect to viral success is evasion from immune clearance. Disruption of either the infection or replication phases should effectively inhibit the SARS-CoV-2 life cycle. Accordingly, many efforts focus on neutralizing interaction of the viral S protein with ACE2 (*4, 46*). Other efforts strive to interfere with the viral life cycle after it invades target cells, and many focus on catalytically active proteins encoded by the SARS-CoV-2 genome (*47*).

Here, we analyzed one small, unstable SARS-CoV-2 protein, ORF9c in the context of an epithelial lung cancer cell line. Limited overlap between published studies of the ORF9c interactome can be attributed to their use of different cell system (HEK293 compared with A549 lung cancer cells used here, as the use of different filtering criteria (*19*). However, many of the cellular changes elicited by ORF9c in our study also occur following infection with full replicative SARS-CoV-2 virus (*22*). Those phenotypes included changes in IFN and other cytokine signaling, immune recognition (including antigen presentation; dendritic cell, T cell, and acute immune responses; and pattern recognition), cell cycle, and the complement system, all of which were downregulated by ORF9c. Additionally, similar to cells infected with the virus or expressing ORF9c, IL-1, IL-6, and p38 MAPK signaling pathways were upregulated. The primary change identified in our analysis was deregulation of the IFN system, coupled with changes in cytokines associated with TNF and STAT signaling and factors implicated in innate immunity. In addition to mediating an antiviral response, aberrant IFN signaling is also critical for numerous pathological indications linked to COVID-19 (*48*). Thus, we concluded that SARS-CoV-2 ORF9c elicits pathologies not seen with previously characterized coronavirus prototypes, primarily through effective modulation of IFN signaling. Our findings suggested that ORF9c enables cells to escape from immune surveillance through by reducing HLA abundance and antigen presentation, while also slowing cell replication, which could viral replication of infected cells.

Strikingly, ORF9c is predicted have a transmembrane domain and we found that the ORF9c interactome was mostly comprised of membrane-associated proteins in multiple organelles, including ER, Golgi, mitochondria, cell surface membrane, and peroxisomes. Indeed, many of the cellular changes that we observed following ORF9c expression are associated with membrane proteins or pathways mediated by proteins that associate with the membranes of various cellular compartments. Importantly, SARS-CoV-2 ORF9c is the first human coronavirus ORF9c protein that has acquired this putative transmembrane sequence. Mutations have been acquired along the course of evolution of ORF9c, although ~80% of the SARS-CoV-2 ORF9c sequence is identical to the ortholog in other coronaviruses, although greater similarity was identified with the bat SARS-CoV-2 sequence. The membranal anchoring capability identified in SARS-CoV-2 ORF9c is novel feature that may mediate the effect on IFN signaling, antigen presentation, and immune evasion phenotypes, characteristics that make SARS-CoV-2 much more virulent and pathogenic than other coronaviruses. Notably, 0.7-1.4% of patients were found to possess a mutation that is expected to impair transmembrane domain of SARS-CoV-2 ORF9c (*19*); awaiting future assessment of clinical outcome, our data would predict a better clinical outcome, distinguishing these patients from those harboring the transmembrane domain. Correspondingly, the interactome for the less virulence and pathogenic SARS-CoV-1 ORF9c (*49*) did not overlap with that for SARS-CoV-2 ORF9c.

A notable signature that we identified is the upregulation of histone and histone deacetylase-related factors, which suggested that histone modification may underlie the transcriptional repression. The increased transcription of AP1 family members (FOSB, FOSL1, CREB, and ATF3), which participate in the cellular stress response, may reflect the response to stress imposed by ORF9c, which, in turn, can limit immune-related signaling identified in our study. Another remarkable signature of ORF9c expression in A459 cells was the association with UBP components. Together with the observations on cellular immune pathways, this association with the UBP suggested that ORF9c induces changes in UBP components that alter the stability of cellular proteins implicated in cytokine signaling, antigen presentation, innate immunity, and the cell cycle. Additionally, we identified UPR components as important for ORF9c instability, suggesting that this protein is misfolded or at least recognized as a misfolded protein by the host cell. In this scenario, we propose that misfolded ORF9c engages UPR (through VCP) and the UBP, which clears this protein. By engaging the UBP, ORF9c promotes enhanced proteasome activity as suggested by our proteome analysis.

Our analysis revealed that interfering with VCP activity blunted the transcriptional repressive effects of SARS-CoV-2 ORF9c on impact immune system components, such as IRF9, INFGR1, ISG15, SOCS1 and TAP1. Proteasome inhibition with bortezomib was also effective, although not as consistently effective as MNS-873, the VCP inhibitor. These findings suggested that inhibition of VCP or the proteasome, which has inhibitors currently in clinical trials for cancer (*50, 51*), may be considered among therapeutic measures to fight SARS-CoV-2 virulence and pathologies. However, we cannot exclude the possibility that ORF9c stability and degradation mechanisms differ based on cell type or the activity of other viral proteins in infected cells. Another potential therapeutic opportunity involves targeting the membrane association of ORF9c, because this is a unique feature of the protein in the SARS-CoV-2 coronavirus. Thus, identifying small molecules that could interfere with ORF9c localization to the membrane could limit ORF9c function and impede the ability of the virus to evade the immune response and reduce viral replication.

Given that ORF9c is expected to affect immune evasion, virulence and pathogenesis, additional studies should assess the consequences of ORF9c inhibition in vivo, using primates and possibly mouse models where SARS-CoV-2 shown to impact IFN signaling and immune response (*48*).

## Materials and Methods

### Cell lines

A549 cells were cultured in Dulbecco’s Modified Eagle’s Medium (Corning) supplemented with 10% Fetal Bovine Serum (Gibco, Life Technologies) and 1% Penicillin/Streptomycin (Corning) and maintained at 37°C in a humidified atmosphere of 5% CO_2_.

### Transfection

Ten million A549 cells were plated in p15 dishes and transfected using Jet Prime (Polyplus) with 15 μg of Strep-tagged expression ORF constructs or empty vector. Samples were harvested 24h after transfection using Dulbecco’s Phosphate Buffered Saline without calcium and magnesium (D-PBS) and supplemented with 10 mM EDTA. Cell pellets were frozen and stored at −80°C. At least three biological replicates were independently prepared for immunoprecipitation or MG132 treatment.

### Immunoblotting

After 24h transfection, A549 were treated with 10μM MG132 (Selleck) during 4h. Whole cell extracts were prepared in RIPA buffer (Thermo Fisher) complemented with cOmplete mini EDTA-free protease and PhosSTOP phosphatase inhibitor cocktails (Roche). Following protein quantification using Pierce’s BCA kit, 5x Laemmli buffer was added and the mix was boiled for 5 minutes. SDS-PAGE resolved proteins were transferred to nitrocellulose membranes and incubated with Strep-tag (Biolegend 688202), β-Tubulin (abcam ab6046) primary antibodies. Secondary antibodies were used at 1:5000.

### Immunoprecipitation

Immunoprecipitation of streptavidin-tagged CoV-2 ORFs was performed as previously described (*19*). Briefly, frozen cell pellets were thawed on ice for 15-20 minutes and suspended in 1ml Lysis Buffer with 50 mM Tris-HCl, pH 7.4 at 4°C, 150 mM NaCl, 1 mM EDTA and supplemented with 0.5% Nonidet P-40 Substitute, Complete mini EDTA-free protease and PhosSTOP phosphatase inhibitor cocktails (Roche). Samples were centrifuged 10 minutes at 4°C at 13,000g. Protein quantification was performed using Pierce’s BCA quantification kit as per the manufacturer’s indications. Supernatants (1 mg protein) were incubated 2h at 4°C with MagStrep “type3” beads (30 μl; IBA Lifesciences) that had been previously equilibrated twice with 1 ml Wash Buffer (IP Buffer supplemented with 0.05% NP40). Beads were washed five times with 1 ml Wash Buffer and then five times with 1 ml Ammonium Bicarbonate 50mM.

### RNA-seq analysis

Raw FASTQ files were processed using cutadapt v1.18 (*52*) to remove adapters. RNA-Seq sequencing reads were aligned using STAR aligner version 2.7 (*53*) based on human genome version 38 and Ensemble gene annotation version 84. Gene expression quantification was performed based on RSEM v1.3.1 (*54*). Gene differential expression was performed using estimated read counts from RSEM by the R Bioconductor package DESeq2 following generalized linear model based on negative binomial distribution (*55*). Genes with Benjamini-Hochberg (BH) corrected p value< 0.05 and fold change >=2 or <=1/2 were selected as significantly differentially expressed genes.

### Pathway and network analysis

Significant differentially expressed genes and proteins were then analyzed using Ingenuity Pathway Analysis (Qiagen, Redwood City, USA) using Canonical Pathways and Upstream Regulators. Canonical Pathway analysis results with BH corrected P<0.1 and Upstream Regulators with P<0.001 were shown in the Supplementary Tables. Differentially expressed genes and differentially expressed proteins were further analyzed using Metascape for MCODE network analysis (*56*). Subcellular locations of proteins were analyzed using DAVID based on Gene Ontology Cellular Component category (*57*) and with the aid of Protein Atlas Protein localization information. Because proteins targeted by ORF9c in interactome and ubiquitinome and proteins targeted by MG132 were usually changed at smaller fold. Differentially expressed genes in Figure 4D, Figure 5A and 5B were selected based on BH correct P <0.05 without fold change cutoff. Differentially expressed proteins in Figure 4D, Figure 5A and 5B were selected based on P <0.01 without fold change cutoff.

### Sequence analysis of ORF9c protein

Protein sequences similar to SARS-CoV-2 ORF9c were retrieved through NCBI Blastp using the nr database (*58*). Sequence alignment was performed using Clustalo (*59*). Phylogenetic tree was built using PhyML algorithm by 100 times bootstrap, and visualized using Seaview (*60*) and Geneious version 2020.2.2 (San Diego, CA). Transmembrane domain prediction was performed using TMHMM web server v2.0 (*38*). Transmembrane domain was predicted based on TMHMM posterior probability more than 0.5.

### Statistical analysis

Statistical test results of RNA-Seq, proteomics and interactome data provided in Supplementary Tables. Analyses of omics data in this study were performed using R customized scripts. Statistical analysis of proteomics data sets was performed using *MSstats* (label-free data) and MStatsTMT (TMT data) bioconductor package. Differential expression of RNA-Seq was performed using DESeq2 bioconductor package following Negative Binomial Distribution and Wald test. Pathway enrichment and upstream regulator analyses were performed using IPA following Fisher’s Exact Test and Z-score calculation considering directional changes in IPA database.

### Data deposits – GSE / public datasets

RNA-Seq data sets were uploaded to NCBI GEO with accession number GSE TBA. Proteomics data were uploaded to ProteinXchange with accession number (TBA). Public datasets used in this study was processed by Coronascape component of Metascape (*56*), including Stukalov *et al*. (*39*) from BioRxiv, and Blanco-Melo *et al*. (*22*).

**Supplementary Materials include Supplemental Methods, Supplemental Figures and Legends (3) and Supplemental Tables (5) are found online at: www.xxx.**

## Acknowledgments

We thank members of the Krogan and Ronai laboratories for help with obtaining reagents and conducting data analysis and for discussions. We also thank Dr. Yingyao Zhou from Genomics Institute of the Novartis Research Foundation for advice for using Metascape for analyses performed in this study. Editorial services provided by Nancy R. Gough (BioSerendipity, LLC, Elkridge, MD)

## Authors contributions

ZAR, RP conceived the research plan; ADA, YF, CCY, performed the experiments; ARC, JY, RM, performed analyses for proteome and transcriptome; HK, AJD, NK, RP, ZAR, reviewed data and wrote the manuscript; DEG provided reagents.

## Data and materials availability

All datasets will be deposited in publicly available data sets prior to publication; all reagents and study protocols are available by requests from the corresponding authors.

## Supplemental Materials

### Supplemental Methods

#### Affinity purification coupled to mass spectrometry

Beads were resuspended in 8M urea, 50 mM ammonium bicarbonate, and cysteine disulfide bonds were reduced with 10 mM tris (2-carboxyethyl) phosphine (TCEP) at 30°C for 60 min. Cysteines were then alkylated with 30 mM iodoacetamide (IAA) in the dark at room temperature for 30 min. Following alkylation, urea was diluted to 1 M urea, and proteins were digested overnight with mass spec grade Trypsin/Lys-C mix (Promega, Madison, WI). Finally, beads were pulled down and the peptide solution collected in a new tube. Affinity purification was carried out in a Bravo AssayMap platform (Agilent) using AssayMap streptavidin cartridges (Agilent). Digested peptides were then desalted in a Bravo AssayMap platform (Agilent) using AssayMap C18 cartridges and dried down in a SpeedVac concentrator.

Prior to LC-MS/MS analysis, dried peptides were reconstituted with 2% ACN, 0.1% FA and concentration was determined using a NanoDrop™ spectrophometer (ThermoFisher). Samples were then analyzed by LC-MS/MS using a Proxeon EASY-nanoLC system (ThermoFisher) coupled to an Orbitrap Fusion Lumos mass spectrometer (Thermo Fisher Scientific). Peptides were separated using an analytical C18 Aurora column (75μm × 250 mm, 1.6 μm particles; IonOpticks) at a flow rate of 300 nL/min (60°C) using a 75-min gradient: 1% to 5% B in 1 min, 6% to 23% B in 44 min, 23% to 34% B in 28 min, and 34% to 48% B in 2 min (A= FA 0.1%; B=80% ACN: 0.1% FA). The mass spectrometer was operated in positive data-dependent acquisition mode. MS1 spectra were measured in the Orbitrap in a mass-to-charge (*m/z*) of 375 – 1500 with a resolution of 60,000 at *m/z* 200. Automatic gain control target was set to 4 × 10^5^ with a maximum injection time of 50 ms. The instrument was set to run in top speed mode with 2-second cycles for the survey and MS/MS scans. After a survey scan, the most abundant precursors (with charge state between +2 and +7) were isolated in the quadrupole with an isolation window of 0.7 *m/z* and fragmented with HCD at 30% normalized collision energy. Fragmented precursors were detected in the ion trap as rapid scan mode with automatic gain control target set to 1 × 10^4^ and a maximum injection time set at 35 ms. The dynamic exclusion was set to 20 seconds with a 10 ppm mass tolerance around the precursor.

Statistical analysis of interactome data was carried out using in-house R script (version 3.5.1, 64-bit), including R Bioconductor packages such as limma and MSstats. First, feature (a peptide sequence with potential amino acid modifications) intensities were log2-transformed and *loess*-normalized within each ORF or control batch to account for systematic errors. Testing for differential abundance was performed using MSstats bioconductor package based on a linear mixed-effects model. Importantly, the log2FC and p-value of proteins missing completely in the negative control (Strep-tagged GFP) were imputed as follows. The imputed log2FC was calculated as the sum of protein intensity (i.e., sum of peptide intensities of a given protein within a given sample) across replicates of the ORF pulldown and divided by 3.3. The imputed p-value was calculated by dividing 0.05 by the number of replicates the protein was confidently identified in the pulldown group. The PPI score is not used for filtering but to indicate significance of different candidates in the list.

High confidence interacting proteins were selected using the following filtering criteria: log2FC > 3.3 (10x) and a p-value <0.025 (to include the p-value of proteins detected in at least 2 ORF9c pulldown replicates but not detected in the negative controls). We also considered the ‘crapomeScore’ < 0.5, which is the fraction of single affinity purification experiments a given protein-interacting candidate receives in the Crapome database (crapome.org). A score of 1 means the candidate is identified in all experiments in that database.

#### Global proteome profiling

Cells were lysed in UAB buffer (8M urea, 50 mM ammonium bicarbonate (ABC) and Benzonase 24U/100ml) with vigorous shaking (20 Hz for 10 min at room temperature using a Retsch MM301 instrument). Lysates were centrifuged at 14,000xg for 10 minutes to remove cellular debris, and protein concentration in supernatants was determined using bicinchoninic acid (BCA) protein assay (Thermo Scientific). Proteins were reduced with 5 mM tris (2-carboxyethyl) phosphine (TCEP) at 30°C for 60 min, and subsequently alkylated with 15 mM iodoacetamide (IAA) in the dark at room temperature for 30 min. Urea was then diluted to 1 M urea using 50 mM ammonium bicarbonate, and proteins digested overnight with mass spec grade Trypsin/Lys-C mix (1:25 enzyme/substrate ratio). Samples then were acidified with formic acid (FA) and desalted using AssayMap C18 cartridges mounted on an Agilent AssayMap BRAVO liquid handling system. Cartridges were sequentially conditioned with 100% acetonitrile (ACN) and 0.1% FA, and samples were loaded and washed with 0.1% FA, and peptides eluted with 60% ACN, 0.1% FA. Finally, organic solvent was removed in a SpeedVac prior to LC-MS/MS analysis.

For TMT sample preparation, total peptide amount was determined using a NanoDrop spectrophotometer (Thermo Scientific), and 25 micrograms of sample was labeled with one TMTpro tag according to the manufacturer’s recommendations. The pooled TMT sample was dried in a SpeedVac, resuspended in 0.1% FA and desalted using a C18 TopTip (PolyLC, Columbia, MD) according to the manufacturer’s recommendations. Finally, organic solvent was removed in a SpeedVac. The dried pooled sample was reconstituted in 20 mM ammonium formate, pH ~10, and fractionated using a Waters Acquity BEH C18 column (2.1× 15 cm, 1.7 μm pore size) mounted on an M-Class Ultra Performance Liquid Chromatography (UPLC) system (Waters). Peptides were separated using a 33-min gradient: 1% to 5% in 0.5 min, 5% to 23.5% B in 1 min, 23.5% to 40% B in 23 min, 40% to 45% B in 1.5 min, 45% to 60% B in 2 min, 60% to 70% B in 4 min, and 70%B to 90% in 1 min (A=20 mM ammonium formate, pH 10; B = 100% ACN). A total of 24 fractions were collected and pooled non-contiguously into 12 fractions (i.e., 1+13, 2+14, 3+15, etc.). Pooled fractions were dried to completeness in a SpeedVac prior to mass spectrometry analysis.

For LC-MS/MS (TMT) analysis, dried peptide fractions were reconstituted with 2% ACN, 0.1% FA and analyzed by LC-MS/MS using a Proxeon EASY nanoLC system (Thermo Fisher Scientific) coupled to an Orbitrap Fusion Lumos mass spectrometer (Thermo Fisher Scientific). Peptides were separated using an analytical C18 Aurora column (75μm × 250 mm, 1.6μm particles; IonOpticks) at a flow rate of 300 μl/min using a 75-min gradient: 1% to 6% B in 1 min, 6% to 23% B in 44 min, 23% to 34% B in 28 min, and 27% to 48% B in 2 min (A= FA 0.1%; B=80% ACN: 0.1% FA). The mass spectrometer was operated in positive data-dependent acquisition mode. MS1 spectra were measured in the Orbitrap with a resolution of 60,000 (AGC target: 4e5; maximum injection time: 50 ms; mass range: from 350 to 1500 m/z). The instrument was set to run in top speed mode with 3 s cycles for the survey and the MS/MS scans. After a survey scan, tandem MS was performed in the *Ion Routing Multipole HCD-Cell* on the most abundant precursors by isolating them in the quadrupole (Isolation window: 0.7 m/z; charge state: + 2-7; collision energy: 35%). Resulting fragments were detected in the Orbitrap at 50,000 resolution (First mass: 110 m/z; AGC target for MS/MS: 1e5; maximum injection time: 105 ms). The dynamic exclusion was set to 20 s with a 10 ppm mass tolerance around the precursor and its isotopes.

For TMT data processing, all mass spectra from were analyzed with MaxQuant software version 1.5.5.1. MS/MS spectra were searched against the *Homo sapiens* Uniprot protein sequence database (downloaded January 2018) and GPM cRAP sequences (commonly known protein contaminants). Reporter ion MS2 type was selected along with TMT 16plex option. Precursor mass tolerance was set to 20ppm and 4.5ppm for the first search, where initial mass recalibration was completed, and the main search, respectively. Product ions were searched with a mass tolerance 0.5 Da. The maximum precursor ion charge state used for searching was 7. Carbamidomethylation of cysteine was searched as a fixed modification, while oxidation of methionine and acetylation of protein N-termini were searched as variable modifications. The enzyme was set to trypsin in a specific mode and a maximum of two missed cleavages was allowed for searching. The target-decoy-based false discovery rate (FDR) filter for spectrum and protein identification was set to 1%.

Statistical analysis of TMT data was carried out using in-house R script (version 3.5.1, 64-bit), including R Bioconductor packages. First, TMT reporter intensities were log2-transformed and normalized (*loess* normalization) across samples to account for systematic errors. Then, all non-razor peptide sequences were removed from the list. Protein-level quantification and statistical testing for differential abundance were performed using *MSstatsTMT* bioconductor package

For LC-MS/MS (label-free) analysis, dried peptides were reconstituted with 2% ACN, 0.1% FA, and concentration was determined using a NanoDrop™ spectrophometer (ThermoFisher). Samples were then analyzed by LC-MS/MS using a Proxeon EASY-nanoLC system (ThermoFisher) coupled to an Orbitrap Fusion Lumos mass spectrometer (Thermo Fisher Scientific). Peptides were separated using an analytical C18 Aurora column (75μm × 250 mm, 1.6 μm particles; IonOpticks) at a flow rate of 300 nL/min (60°C) using a 75-min gradient: 1% to 5% B in 1 min, 6% to 23% B in 44 min, 23% to 34% B in 28 min, and 34% to 48% B in 2 min (A= FA 0.1%; B=80% ACN: 0.1% FA). The mass spectrometer was operated in positive data-dependent acquisition mode. MS1 spectra were measured in the Orbitrap in a mass-to-charge (*m/z*) of 375 – 1500 with a resolution of 60,000 at *m/z* 200. Automatic gain control target was set to 4 × 10^5^ with a maximum injection time of 50 ms. The instrument was set to run in top speed mode with 2-second cycles for the survey and the MS/MS scans. After a survey scan, the most abundant precursors (with charge state between +2 and +7) were isolated in the quadrupole with an isolation window of 1.6 *m/z* and fragmented with HCD at 30% normalized collision energy. Fragmented precursors were detected in the ion trap as rapid scan mode with automatic gain control target set to 1 × 10^4^ and a maximum injection time set at 35 ms. The dynamic exclusion was set to 20 seconds with a 10 ppm mass tolerance around the precursor.

For processing label-free LC-MS/MS data, all raw files were processed with MaxQuant (version 1.5.5.1) using the integrated Andromeda Search engine against a target/decoy version of the curated human Uniprot proteome without isoforms (downloaded in January of 2020) and the GPM cRAP sequences (commonly known protein contaminants). First search peptide tolerance was set to 20 ppm, and main search peptide tolerance was set to 4.5 ppm. Fragment mass tolerance was set to 20 ppm. Trypsin was set as the enzyme in specific mode, and up to two missed cleavages was allowed. Carbamidomethylation of cysteine was specified as fixed modification and protein N-terminal acetylation and oxidation of methionine were considered variable modifications. In addition, the phosphopeptide-enriched samples were also searched with phosphorylation of serine, threonine or tyrosine considered as variable modification. The target-decoy-based false discovery rate (FDR) filter for spectrum and protein identification was set to 1%.

Statistical analysis of label-free proteomics data was carried out using in-house R script (version 3.5.1, 64-bit), including R Bioconductor packages. First, peptide feature intensities (MaxQuant evidence table) were log2-transformed and normalized (*loess* normalization) across samples to account for systematic errors. Then all non-razor peptide sequences were removed from the list. Protein-level quantification and statistical testing for differential abundance were performed using *MSstats* bioconductor package.

#### siRNA screen

The 1141 Ubiquitome siRNA library was generated from the ON-TARGETplus® SMARTpool® human siRNA genome library (a pool of 4 unique siRNAs, Dharmacon, Thermo Scientific). siRNA was used at a final concentration of 10 nM with 0.01μl Lipofectamine RNAiMAX (Invitrogen) reagent and 600 cells in a final volume of 50μl/well in black and clear bottom CellCarrier-384 well plates (PerkinElmer). siRNAs were pre-spotted on a Bravo automated liquid handling platform (Agilent), and all other liquid additions performed with a Thermo/Matrix Wellmate. Plates were incubated at 37°C in 10% CO_2_ for 72h. For positive controls, MG132 was added to wells 4 hours before fixation at a final concentration of 10uM. Cells were then fixed (paraformaldehyde 3.7% in PBS, 10min), permeabilized (0.5% Triton X-100 in PBS, 5 min), blocked (3% BSA in PBS, 1 h), and then incubated first with primary antibody (anti-Strep-tag, Thermo Fisher, cat# MA5-17282,1:1000 in 3% BSA) and then with secondary antibody(Alexa Fluor 488 Donkey-anti mouse antibody, Invitrogen, 1:1000 in 3% BSA). Wash buffers were PBS plus 0.1% Triton X-100. siRNAs pools containing 48 non-targeting siRNAs (Dharmacon) were used as a negative control.

#### High-content imaging analysis

Cells were imaged with an IC200 high-content screening system (Vala Sciences) using a 20X objective to visualize Strep-ORF9c proteins (Alexa 488) and nuclei (DAPI). Four images were obtained from different fields in each well for 384-well plates. Images were analyzed with Acapella high-content imaging and analysis software for valid cell numbers per field and to determine average Alexa 488 intensity per cell. A549 cells expressing SARS-CoV-2 Strep-ORF9c were treated with DMSO or MG132 (10uM) served as negative and positive imaging controls, respectively. Plate-to-plate variability was normalized using a control-based method; associated control samples were aggregated, and the mean and variance across wells were determined. The Alexa488 mean intensity for all wells with siRNA knockdown was normalized using unique non-targeting siRNAs included in each plate as reference data points. The top 36 scoring hits were obtained using a threshold of > 1.46-fold increase in average intensity from duplicates (p-value <0.05). Ten of the 36 siRNA pools were selected for confirmation in a secondary deconvolution screen. For that screen, quantification data were converted to a Z-score, and the average Z-score from data in triplicate plates was determined. Genes were defined as confirmed screen hits if they had 3 or more individual positive siRNA score (cut-off of >3 SD).

#### qPCR and primers

A549 cells were transfected with CoV-2 ORFs plasmids and samples harvested at 24h, 48 and 72 h. Total RNA was extracted using RNAeasy (Qiagen) and transcribed into cDNA by cDNA Archive Kit (Applied Biosystems) according to the manufacturer′s instructions. Expression of CoV-2 ORFs accessory protein transcripts was analyzed by quantitative real-time PCR (qRT-PCR) using the following primers: IFNGR1 F AGCAGGAAGTCGATTATGATCCC, R CTGGCACTGAATCTCGTCACA; ISG15 F CGCAGATCACCCAGAAGATCG, R TTCGTCGCATTTGTCCACCA; PSMB8 F GGTCCTACATTAGTGCCTTACGG, R CGCAGATAGTACAGCCTGCATT; SOCS1 F TTTTCGCCCTTAGCGTGAAGA, R GAGGCAGTCGAAGCTCTCG; TAP1 F GCAAGACGACTTACTCTGGGT, R GGATCTGACACCACTGGACC. Cycle threshold values were determined and normalized to the housekeeping gene GAPDH for each experiment. Relative gene expression was calculated by the 2^−ΔΔ*C*t^ method. Results are expressed as means ± SDs of 3 independent experiments. Statistical analysis was performed using Student′s t-test. A *p*-value of < 0.05 was considered statistically significant.

### Supplementary Figures Legend

**Figure S1.**
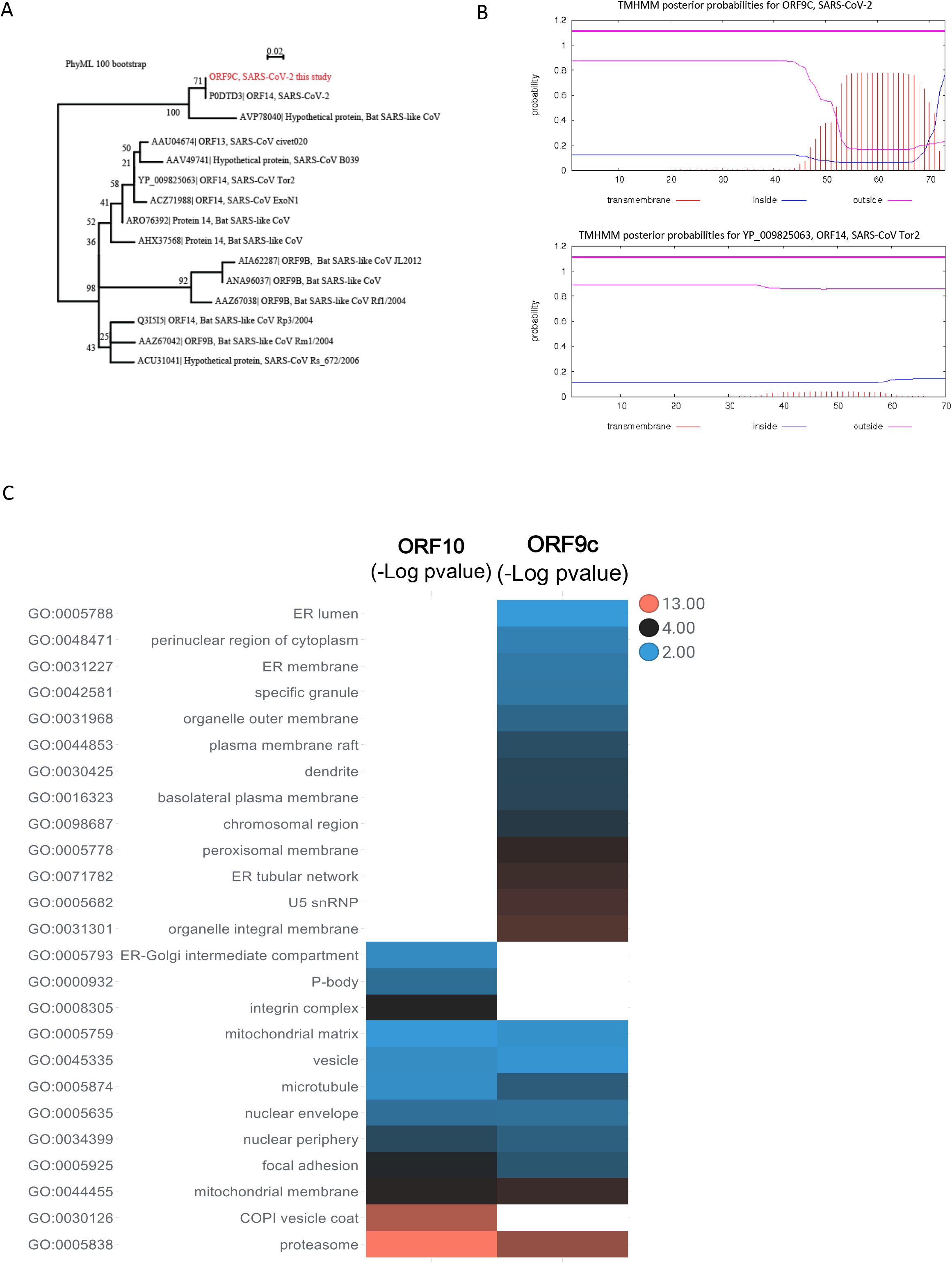
Phylogenetic analysis and transmembrane prediction for SARS-CoV-2 ORF9c. **A**. Phylogenetic tree of SARS-CoV-2 ORF9c in indicated strains of coronavirus **B**. (Upper panel) TMHMM prediction that the SARS-CoV-2 ORF9c C-terminal contains a transmembrane domain spanning ~20 amino acids. (Lower panel) Comparable analysis of SARS-CoV ORF14 indicating a low probability of a transmembrane domain. **C.** Gene Ontology (GO) cellular component enrichment analysis of strep-tagged ORF9c- and strep-tagged ORF10-interacting proteins in A549 cells detected by APMS revealed an enrichment of membrane or membrane-related terms in ORF9c. The heatmap shows the −log p-value of enrichment analysis performed in Metascape.

**Figure S2.**
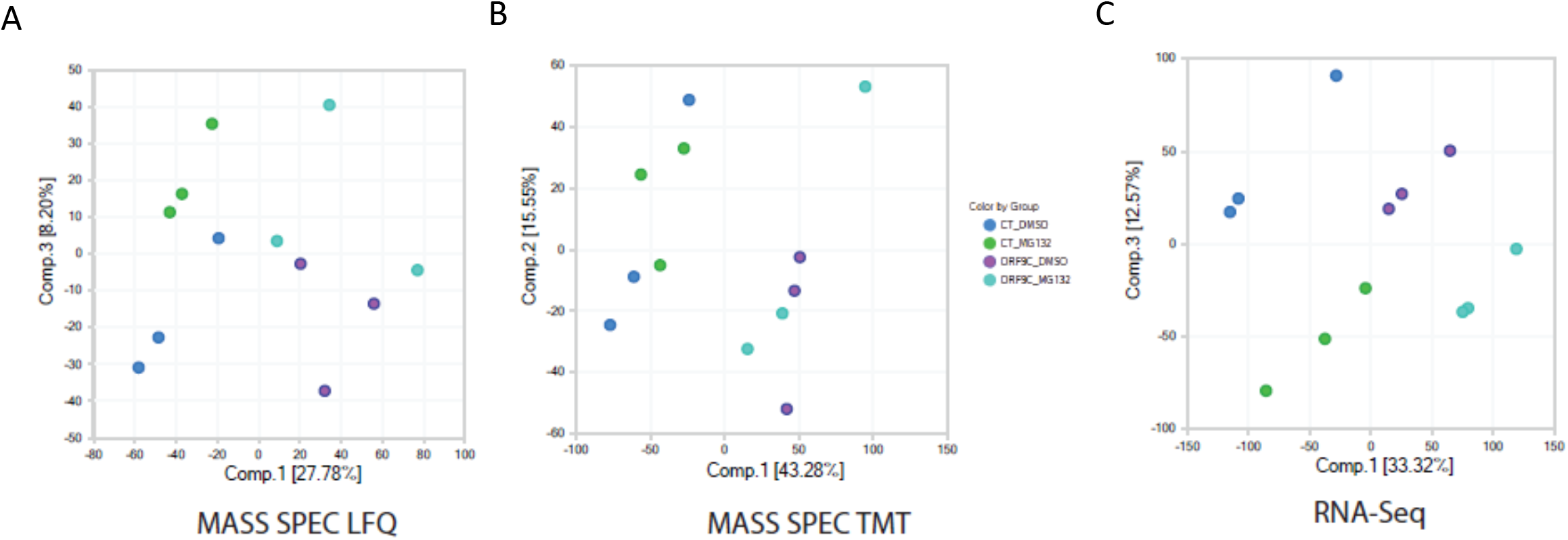
Principal component analysis of the proteomic and transcriptomic data for A549 cells expressing SARS-CoV-2 ORF9c. **A, B**. PCA of MS/LFQ and MS/TMT data showing ORF9c expression as a major determinant, contributing to 28% to 43% of respective sample variance. MG132 treatment also distinguished samples by contributing to 15 – 20% of total variance. **C**. PCA showing ORF9c expression as the major factor contributing to 33% of sample variance. MG132 treatment also distinguished samples by contributing to 13% of total variance.

**Figure S3.**
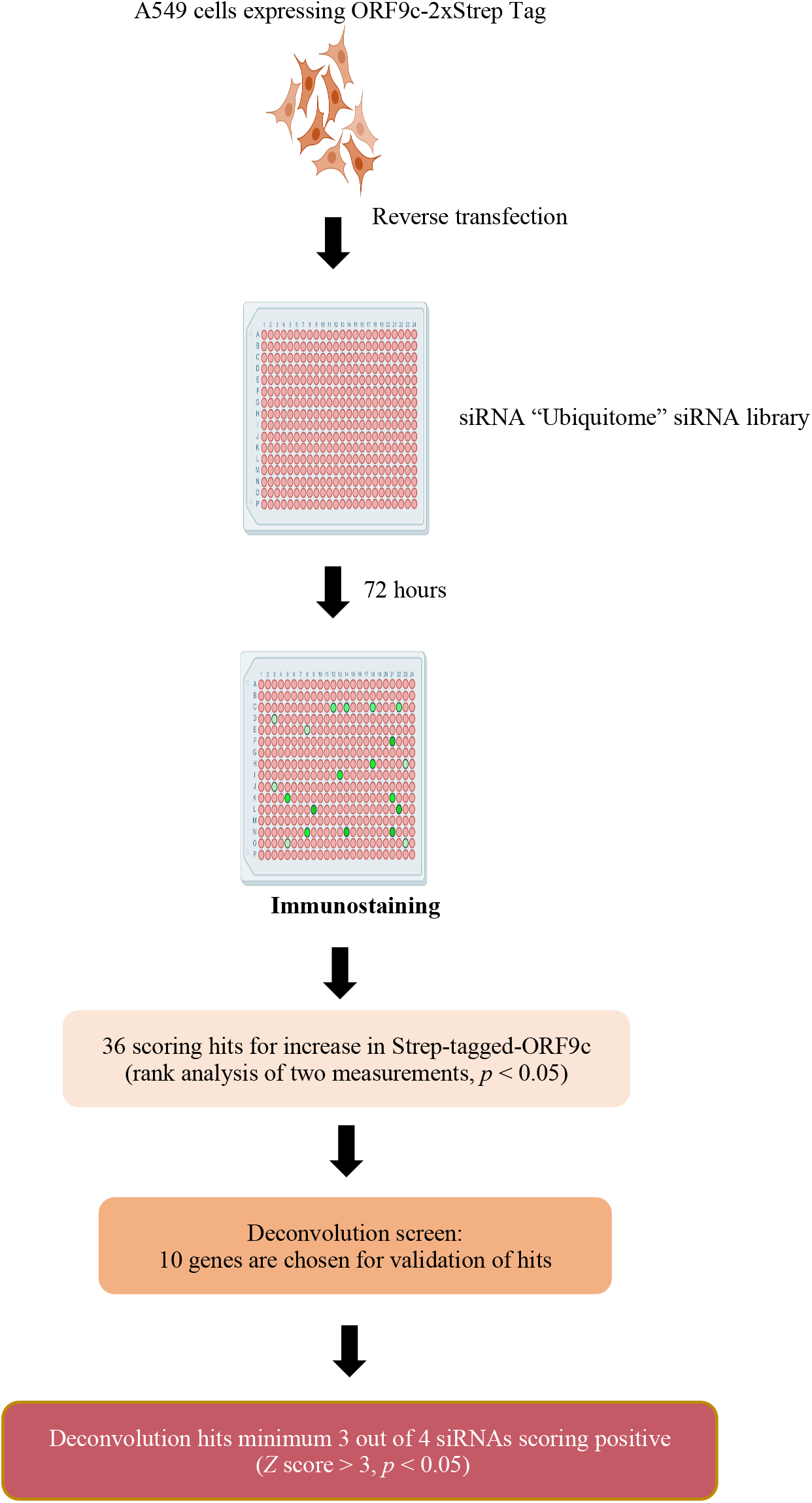
Schematic outline for the siRNA screen performed to identify UBP components that can halt ORF9c degradation. Outlined is the approach we used for siRNA screen of A549 cells that stably express strep-ORF9c using reverse transfection of over 1,100 genes from the UBP library. Microscopy-based screen enabled the identification of siRNA of select UBP components that prevented the degradation of ORF9c.

### Supplemental Tables Legend

**Table S1.** Interactome and proteome data analysis, pathway enrichment and upstream regulators analyses

**Table S2.** Transcriptome data analysis, pathway enrichment and upstream regulators analyses

**Table S3.** Canonical pathway comparison of different omics technologies and public data sets

**Table S4.** Canonical pathway analysis of MG132 reversed genes and proteins

